# A Standardized Ontology for Naming tRNA-derived RNAs Based on Molecular Origin

**DOI:** 10.1101/2022.05.06.490965

**Authors:** Andrew D. Holmes, Patricia P. Chan, Qi Chen, Pavel Ivanov, Laurence Drouard, Norbert Polacek, Mark A. Kay, Todd M. Lowe

## Abstract

Small RNAs processed from transfer RNA are recognized to have regulatory functions distinct from protein synthesis. This rapidly advancing field has led to a constellation of transcript nomenclatures. Building upon the accepted tRNA naming system and linking tRNA-derived small RNAs to their molecular sources, we propose an improved nomenclature. Facilitated by the tDRnamer system, it will promote inter-study comparisons leveraging a biologically-rooted nomenclature for this emerging class of intriguing regulators.

## Main

Transfer RNA, well-studied as the key adapter molecule in protein translation, is now known to be processed into a large collection of smaller RNAs which vary across cell types and species. High-throughput small RNA sequencing methods, including those specialized for tRNAs such as ARM-Seq^1^, Pandora-seq^2^, and others, have led to the identification of tRNA-derived small RNAs (tDRs) in eukaryotes, bacteria, and archaea^3–5^. Functional studies have revealed a plethora of regulatory roles associated with these small noncoding RNAs, including modulation of ribosome biogenesis, protein synthesis, gene silencing, stress response, and plant nodulation^6–9^. Moreover, differential abundance of tDRs observed across tissues and cellular conditions suggests the possibility of leveraging tDRs as biomarkers for early diagnosis of cancer, heart disease, neurological disorders, and other diseases^10^. These small RNAs have been most commonly referred to as “tRNA fragments”, and are often simply categorized by their positions in source tRNAs, namely, 5′ or 3′ fragments, 5′ or 3′ halves, internal fragments, 5′ leaders, or 3′ trailers^4,11^. Contributions from many different research groups have led to a constellation of diverse yet often overlapping names and acronyms, for example, tRFs, tsRNAs, tiRNAs, SHOT-RNAs, tRNA halves, among others. The use of the term “tRNA fragment” may also cause confusion in distinguishing between stable, functional tRNA-derived RNAs, and passive, presumed non-functional degradation products. Although naming algorithms such as License Plates^12^ and databases such as tRFdb^3^ assign systematic identifiers, for various reasons, these names have not been widely adopted in the same way as systematic tRNA gene names. Here we propose a consistent, uniform naming system (Fig. 1a) that is biologically informative and builds upon an existing, proven naming scheme. To make the naming system readily accessible to the research community, we have also developed tDRnamer (Fig. 1b, Supplementary Table 1), a new resource with both web-based and standalone software versions to enable user-friendly information-rich interfaces as well as efficient bulk processing. Given one or more sequences, the tool deterministically generates standardized tDR names; conversely, given one or more previously generated tDR names, it provides exact sequences that avoid any ambiguity while preserving a consistent, descriptive nomenclature. Additional annotations and graphical visualizations produced by tDRnamer further highlight the relatedness of similar tDRs with their source tRNA(s).

**Fig. 1.**
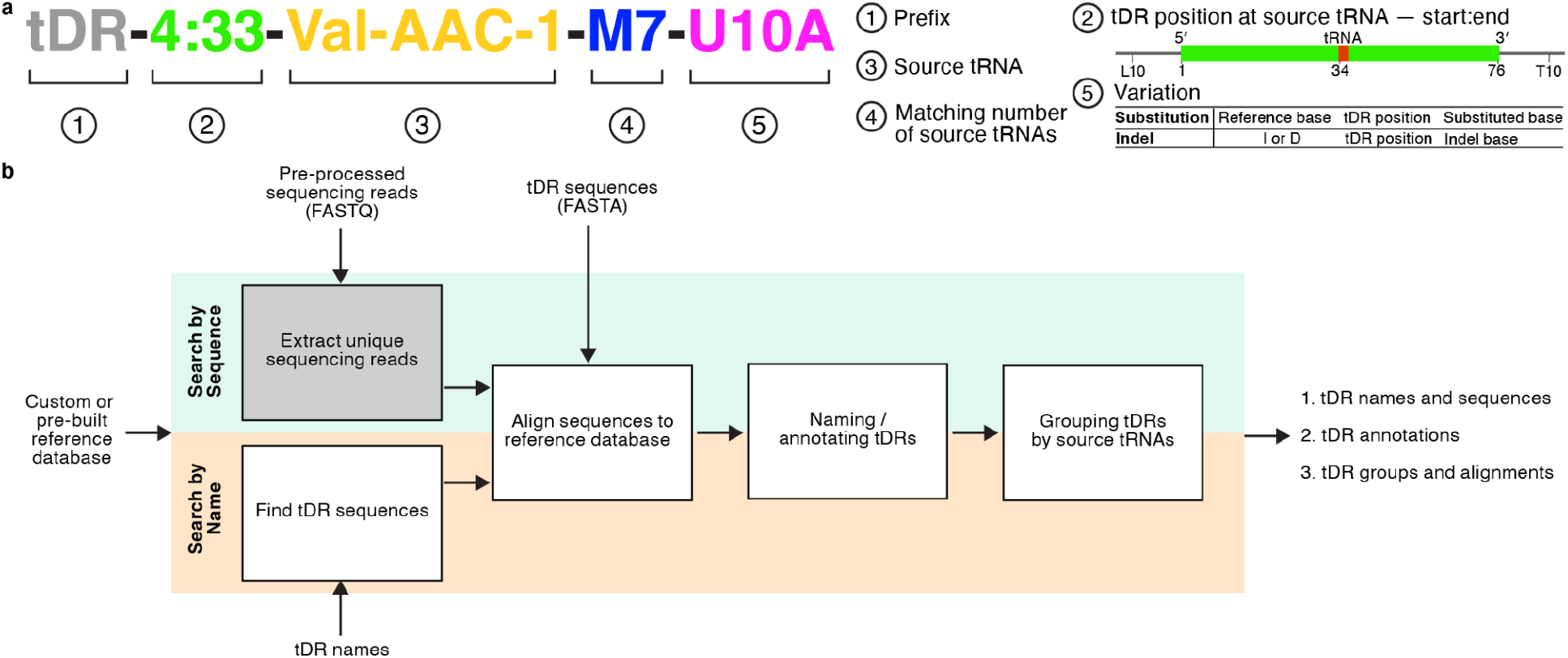
Standardized naming for tRNA-derived RNAs (tDRs) and overview of tDRnamer workflow. **a**. The standardized tDR name contains five components: (1) the prefix that indicates a tDR, (2) position of the tDR relative to its source tRNA in Sprinzl numbering^13^ except for 5′ leader and 3′ trailer positions which precede positions with “L” or “T” respectively, (3) the source tRNA name from GtRNAdb^14^, (4) the number of total matching source tRNA transcripts when more than one exist, and (5) nucleotide variation if it exists between the source tRNA and tDR. **b**. Both the standalone and web server versions of tDRnamer generate tDR names according to the proposed convention, provide tDR annotations, and perform tDR group alignments from input sequences. They also search for tDR sequences with annotations and alignments based on query tDR names. Gray box represents a processing step only available in the standalone version.

The tDR naming system (Fig. 1a) consists of three required components, plus up to two supplemental notations in special cases. First, each name starts with the neutral prefix “tDR” which is general to any functional role. For tDRs that are derived from mitochondrial or plastid tRNAs, “mtDR” and “ptDR” are used respectively. Second, the tDR’s precise start and end positions relative to its source tRNA are specified using the standardized Sprinzl tRNA position numbering^13^ (e.g., in Fig. 1a, “4:33”), which is applicable to all tRNAs, placing the molecule in the rich structural and functional context of over fifty years of tRNA research. This gives the stems and loops consistent base numbers across all tRNAs, including key features such as the anticodon (i.e., positions 34-36). Third, the identifier of the most likely source tRNA found within the Genomic tRNA Database (GtRNAdb)^14^ is appended to the tDR name (e.g., “Val-AAC-1”). Because some regions of different tRNAs in the same species may be identical, tDRs may be derived from multiple indistinguishable parent tRNAs (for example, see Fig. 2c: 5011a exactly matches the 5’ end of 8 different tRNAs). In this case, a fourth component is added to the tDR name to indicate when there is more than one potential source tRNA (e.g., “M7”). Finally, a fifth component of the name may be needed to represent any nucleotide differences between the tDR and the most similar source tRNA. Differences between tDRs and the reference tRNA set could potentially be due to nucleotide misincorporation at certain RNA modifications (common during the reverse transcription step of tRNA sequencing), or natural genetic variation. To describe these tDRs succinctly, a name component is added, consisting of the original and substituted bases (or insertions/deletions) with their relative positions (e.g., “U10A”).

**Fig. 2.**
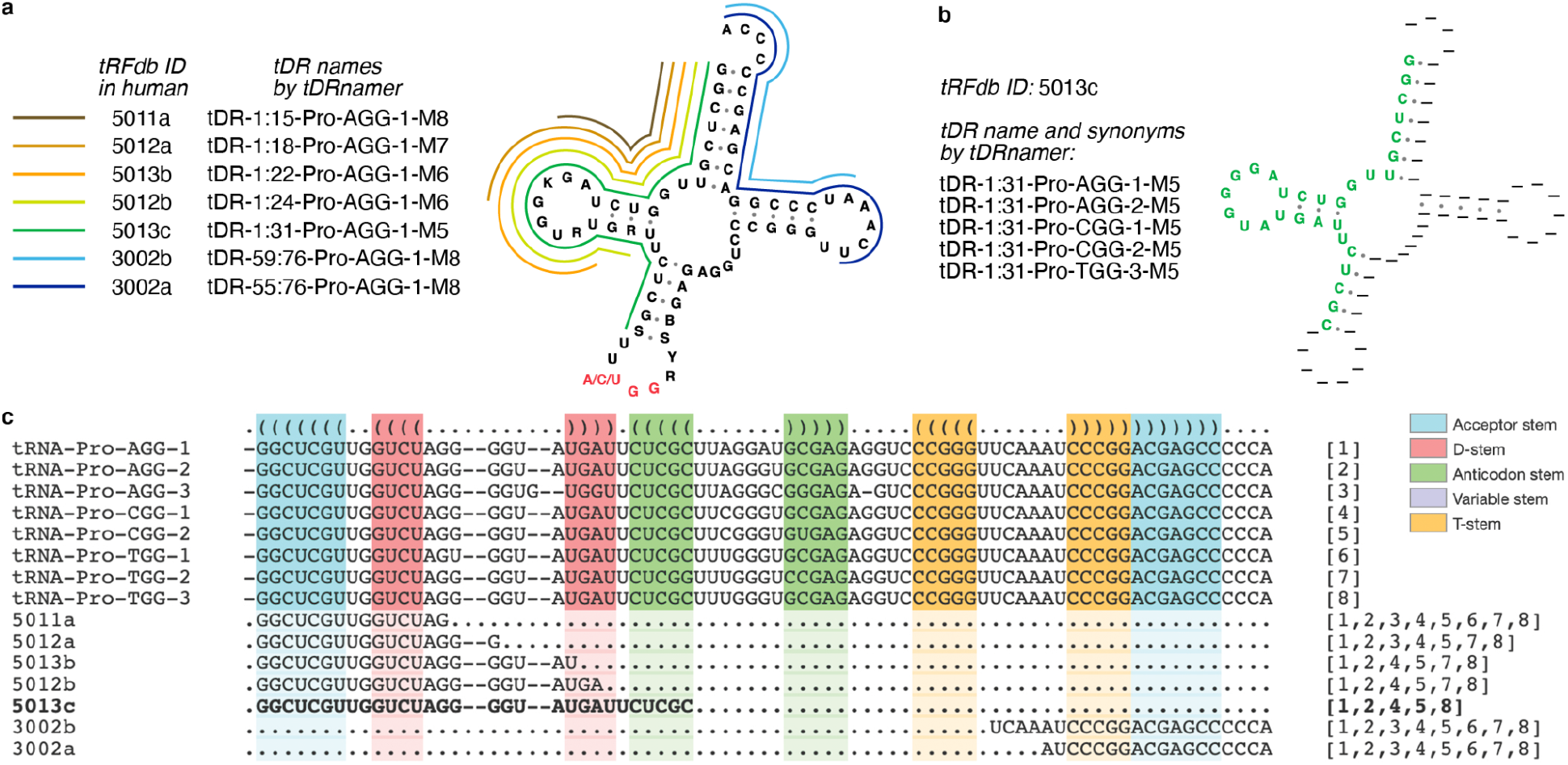
Seven tDRs potentially derived from up to eight human tRNA^Pro^ transcripts. **a**. The tDR names for seven tRFdb entries with tRNA^Pro(AGG)^ as the source tRNA are listed. Colored lines around the consensus tRNA^Pro^ secondary structure represent the derived tDRs. **b**. An example tDR, tRFdb ID 5013c, is located at the 5′ end of tRNA^Pro^ from position 1 to 31 as illustrated in the superimposed secondary structure generated by tDRnamer. The tDR name and synonyms show that its source tRNAs may include two different tRNA^Pro(AGG)^, two different tRNA^Pro(CGG)^, and/or one tRNA^Pro(UGG)^. **c**. tDRs are grouped and aligned with mature tRNA^Pro^ by tDRnamer. The color blocks represent the stems in tRNA secondary structure. The bracketed numbers at the end of each tDR (e.g., [1,2,4,5,8] for 5013c) refer to the five possible source tRNAs shown at the top of the alignment.

To demonstrate application of the new naming system with tDRnamer, we created systematic names for the existing sequence entries in tRFdb^3^, a widely-used, first-of-its-kind database containing sequences from eight different species. Focusing on the human subset as an example, one can glean a wealth of information from examining the tDR names and new analytics, generated in seconds, from the tDRnamer website service. For example, 79% (123 out of 155) of these tDRs are derived from mature tRNAs; the rest are derived from pre-tRNAs, consistent with the tRF type classification (tRF-5, tRF-3, and tRF-1) from tRFdb (Supplementary Data 1). In terms of determining all the potential sources of the tRFdb human entries, tDRnamer found that more than half (88 out of 155) could be derived from more than one source tRNA (i.e., those with “M” in the fourth component of the tDR name), with every possibility given (Supplementary Data 1). tDRnamer also provides multiple-sequence alignments to illustrate the relationships among the set of submitted tDR sequences and all potential source tRNAs. An interesting case involves a group of seven tRFdb entries potentially derived from multiple tRNA^Pro^ with different anticodons (Fig. 2a). For tDR-1:31-Pro-AGG-1-M5 (tRFdb ID: 5013c), the possible synonyms include all five matching source tRNAs (Fig. 2b-c; Supplementary Table 2). In comparison, this information is necessarily absent in the tRFdb IDs^3^ and License Plates^12^ nomenclature. Five of the tDRs align to the 5′ end of the source tRNAs from position 1 as described in names (“1:15”, “1:18”, “1:22”, “1:24”, and “1:31”) while the remaining two align to the 3′ end including the CCA tail ending at position 76 (“59:76” and “55:76”) (Fig. 2a). The group alignments produced by tDRnamer also show that the longest 30-nt tDR (5013c) exactly matches five tRNA^Pro^ and the others match six (5012b, 5013b), seven (5012a) or eight source tRNAs (5011a, 3002a, 3002b) (Fig. 2c). Several other examples of tDR complexity, clarified via alignments and other visualizations produced by tDRnamer, showcase the capabilities of the web analysis interface (Supplementary Figs. 1-2). For instance, three different regions of human tRNA-Gly-GCC-1 are processed into different tDRs of multiple lengths (Supplementary Fig. 1), whereas a tDR derived from 3′ trailer sequence of tRNA-Arg-ACG-1-3 includes multiple single-nucleotide variants (Supplementary Fig. 2). These collective examples show the complexity, breadth, and descriptive capabilities of the proposed tDR naming system. tDRnamer provides the seamless engine for translating RNA sequences to and from the new vocabulary. The interactive web server also should be an indispensable analytic tool to recognize biological relatedness among tDRs and their sources, facilitating advancements in this fast-growing field where comparisons of findings between studies has been unnecessarily difficult.

## Methods

### tDRnamer software design and requirements

tDRnamer is a software package that generates standardized transcript names and annotations for tRNA-derived RNAs (tDRs) and searches tDR sequences by name (Fig. 1b, Supplementary Table 1). The standalone version runs natively as a command-line tool within the Unix/Linux environment, and can also be used via a Docker container or Conda environment in a broader context. The software was developed using Perl and Python for the framework and data processing. The package also uses external bioinformatics tools and data sources as described below in more detail. Researchers who do not have the resources or expertise to work with a command-line tool can use the web server version, developed using HTML, Javascript, and CGI Perl for the execution of the tDRnamer processing engine. The command-line software was designed to be used with transcript sequences in FASTA file format as well as sequencing data in FASTQ format generated by the Illumina sequencing platform. Due to the relatively large file size and much longer processing time for sequencing data, the web server version is limited to support only transcript sequences as input. For the same reason, we recommend that researchers use tDRnamer on a computing server with multi-core processors instead of personal computers when processing sequencing data.

### Reference database generation

tDRnamer requires the use of a reference database that is specifically built with mature and precursor tRNAs of a desired genome. A tool called create_tDRnamer_db is included in the tDRnamer package with which researchers can create the database via tRNAscan-SE 2.0^15^ annotations and sequences downloaded from the Genomic tRNA Database^14^, and the genome sequence of the organism. As a default feature, the tool will exclude pseudogenes and tRNA genes with unknown isotype in the database generation process. Moreover, only the high confidence tRNA set, and genes with consistent isotype model and anticodon, prediction score at least 50 bits, and isotype model score at least 80 bits are included for large eukaryotes to avoid confusion and likely inaccurate associations with tRNA-derived repetitive elements which very rarely produce stable mature tRNAs. tRNA genes removed by this filtering step are outputted in the tRNAscan-SE^15^ output file format (Supplementary Table 3). Researchers can choose to skip this filtering step by using the --skipfilter option or specify alternative score cutoffs with --score and --isoscore options to fit their needs.

The database building tool generates a Bowtie2^16^ index (Supplementary Table 3) that consists of unique mature tRNA and precursor tRNA sequences based on the provided data. The mature tRNA sequences are created with 3′ CCA tails, removal of introns, and addition of the histidine tRNA-specific post-transcriptional 5′ G base at position −1. Unlike eukaryotic tRNAs that always have the 3′ CCA added enzymatically, some bacterial and archaeal tRNAs have genetically encoded 3′ CCA^17^ tails. Therefore, the tool will assess the existence of 3′ CCA in each tRNA gene when the sequence source (--source option) is set for bacteria or archaea, and only make the addition as appropriate. As an additional step, 20 “N” bases flanking each side of the tRNA sequences are added to allow for extra bases appearing off the end of sequenced transcripts, such as potential “CCACCA” ends. Precursor tRNA sequences are created with 50 nt flanking regions to accommodate tDRs that are derived from 5′ leaders or 3′ trailers. When the intergenic region that flanks a tRNA gene is shorter than 50 nt, such as those transcribed in polycistronic operons, only the annotated intergenic region will be included. The reference database also contains the alignments of both mature and precursor tRNA sequences in Stockholm^18^ file format for determining the derived location of tDRs (Supplementary Table 3). These alignments that include both primary sequence and secondary structure information are generated using the cmalign program from Infernal v1.1^18^ with options “--nonbanded --notrunc -g” and clade-specific tRNA covariance models corresponding to the specified sequence source. For alignments of precursor tRNAs, covariance models trained with eukaryotic, bacterial, or archaeal tRNA genes from tRNAscan-SE 2.0^15^ are used. Because tRNAscan-SE covariance models were trained with tRNA genes (some which include introns), the training data were converted to mature tRNA sequences as described above for generating new covariance models using cmbuild from Infernal^18^, with options “--hand --wnone --enone”.

Reference databases for the following model organisms in Eukaryota, Bacteria, and Archaea have been pre-built for the web server version and are available for download: *Arabidopsis thaliana* (TAIR10), *Caenorhabditis elegans* (WBcel235/ce11), *Drosophila melanogaster* (Release 6 plus ISO1 MT/dm6), *Homo sapiens* (GRCh37/hg19 and GRCh38/hg38), *Mus musculus* (GRCm38/mm10 and GRCm39/mm39), *Rattus norvegicus* (Rnor_6.0/rn6 and mRatBN7.2/rn7), *Saccharomyces cerevisiae* S288c, *Schizosaccharomyces pombe* 972h-, *Bacillus subtilis* 168, *Escherichia coli* K-12 MG1655, *Helicobacter pylori* J99, *Mycobacterium tuberculosis* H37Rv, *Haloferax volcanii* DS2, *Methanosarcina barkeri* str Fusaro, *Methanococcus maripaludis* S2, and *Thermococcus kodakarensis* KOD1. Reference databases for additional species may also be requested for use on the web server resource.

### Processing small RNA sequencing reads

When searching by sequence (option --mode as “seq”), tDRnamer automatically determines the input data type by file extension. Small RNA sequencing reads should be contained in a FASTQ file called *.fq or *.fastq, or in Gzip compressed format. The sequencing reads are expected to have been pre-processed, with sequencing adapters and low-quality reads removed, before executing tDRnamer. If paired-end reads are used, the two read ends must be merged into pseudo-single-end reads. Although tDRnamer does not include the pre-processing step, trimadapters.py in the companion tRNA-sequencing data analysis pipeline, tRAX (http://trna.ucsc.edu/tRAX/), can be used for this task. To identify possible tDRs, unique sequencing reads are extracted from the input data as the first step of tDRnamer. Only those reads that have at least ten copies are selected by default, however a different threshold can be specified with --minread option. A sequence file in FASTA format is generated with the identical read count as the suffix of each sequence name (Supplementary Table 4) and can be further processed using the same routine as using a FASTA file for tDRnamer’s input data.

### Identifying tDRs by alignments to reference database

Sequences provided in a FASTA file or as unique sequencing reads extracted from a FASTQ file are aligned to the pre-built reference database using Bowtie2^16^ in the very-sensitive mode which ignores quality scores and allows a maximum of 100 alignments per sequence (--very-sensitive --ignore-quals --np 5 -k 100). Because RNA modifications and editing may exist in tDRs, tDRnamer does not use a hard mismatch threshold to filter alignments. Instead, only all best mappings (highest alignment score – AS tag in SAM file) of the sequence alignments are included. When the sequence source (--source option) is set to eukaryotes, sequences aligned to mature tRNAs and pre-tRNAs are both assessed. Otherwise, only those aligned to mature tRNAs are assessed. If a sequence aligns equally well to a mature tRNA and pre-tRNA, the mature tRNA alignment is preferred and the pre-tRNA alignment is excluded. Using this method, sequences that contain a region flanking the tRNA loci in the genome will be considered to be part of the pre-tRNAs, while sequences that contain no flanking sequence will be considered as derived from mature tRNAs. Sequences that are successfully aligned to tRNAs are further processed as identified tDRs.

### Naming and annotating tDRs

To calculate the Sprinzl^13^ positions and secondary structure regions of tDRs, they are structurally aligned to the reference database of tRNAs. Nucleotide differences including substitutions, insertions, and deletions between tDRs and the matching source tRNAs are determined from the alignments. Using the source tRNA identity information, the starting and ending Sprinzl^13^ positions where the tDR overlaps the source tRNA, and nucleotide differences if they exist, tDRnamer then generates the standardized name for each individual tDR (Fig.1a). If the tDR is derived from multiple source tRNAs, synonyms following the same naming convention are created based on all the identified source tRNAs. In addition to outputting a tDR sequence file in FASTA format, a tab-delimited file containing the tDR names, annotations, and sequences is generated, as well as a file containing the alignments of tDRs with source tRNAs in Stockholm^18^ format (Supplementary Table 4). Furthermore, the tDRnamer web server version uses NAVIEW^19^ to superimpose tDR sequence onto the secondary structure image of its source tRNA for visualization and file download in PNG and postscript formats (Fig. 2, Supplementary Table 4). tDR secondary structure based on minimum free energy is computed using RNAFold^20^ with images rendered by Forna^21^ and VARNA^22^ (Supplementary Fig. 2, Supplementary Table 4).

### Grouping tDRs by source tRNAs

If tDRs identified in the same tDRnamer analysis run are derived from the same source tRNAs, they are grouped together to visualize all tDR products processed from the same parent molecule(s). tDRs that are derived from mature tRNAs are grouped separately from those derived from pre-tRNAs. Alignment of tDRs with their corresponding source tRNAs is based on both primary sequence and secondary structure. Numeric IDs are assigned to the source tRNAs belonging to each group in sequential order of the tRNA transcript name or GtRNAdb gene symbol. tDRs in the alignments are marked with the numeric IDs corresponding to their source tRNA(s). A text file is generated as output which contains the tDR groups, each with the member tDR list, the source tRNAs, the tRNA isotype, nucleotide substitutions (mismatches) if they exist, and the tDR alignments in Stockholm^18^ format (Supplementary Table 4).

### Finding tDR sequences by name

tDR names provided as queries to find their corresponding sequences are split into the defined components according to the naming convention (Fig. 1a) and checked for validity. The source tRNA name and Sprinzl^13^ positions extracted from each valid tDR name are used to search for the corresponding sequence in the reference database. If nucleotide differences between tDR and its source tRNA are encoded in the name, the tDR sequence retrieved from the database will be modified to include the changes. A sequence file in FASTA format is generated as output (Supplementary Table 4). tDR sequences found in the search process are then processed using the same method as searching by sequences described above to obtain the same full annotation and grouping output files (Fig. 1b, Supplementary Table 3).

### Re-annotation of tDRs in tRFdb

tDR data including sequences, mapped positions, and source tRNAs were downloaded from tRFdb^3^ in CSV file format. Sequences were converted into a FASTA file for each genome to be analyzed. tDRnamer reference databases were generated with default options using tRNA annotations from GtRNAdb^14^ release 19 for human assembly GRCh38/hg38, mouse assembly GRCm39/mm39, *Caenorhabditis elegans* assembly WBcel235/ce11, *Drosophila melanogaster* assembly BDGP Rel. 6/dm6, *Schizosaccharomyces pombe* 972h-, *Xenopus tropicalis* assembly Xenopus_tropicalis_v9.1/xenTro9, zebrafish assembly GRCz11/danRer11, and *Rhodobacter sphaeroides* ATCC 17025. tDR sequences in each genome were then analyzed and annotated using tDRnamer with --minlen option set as 13 to allow inclusion of shorter transcripts.

## Data availability

The tDRnamer web server is available at http://trna.ucsc.edu/tDRnamer/. Standalone software can be obtained from GitHub at https://github.com/UCSC-LoweLab/tDRnamer and Docker image at https://hub.docker.com/r/ucsclowelab/tdrnamer. Example data can be downloaded from http://trna.ucsc.edu/tDRnamer/data/examples/. Pre-built reference databases for model organisms are available at http://trna.ucsc.edu/tDRnamer/docs/refdb/. The complete set of tRFdb^3^ data re-annotations by tDRnamer is available at http://trna.ucsc.edu/tDRnamer/data/tRFdb_reannotations/.

## Code availability

The source code of tDRnamer and its web server is available at https://github.com/UCSC-LoweLab/tDRnamer and https://github.com/UCSC-LoweLab/tDRnamer-web respectively, both under the GNU General Public License v3.0.

## Acknowledgements

This work was supported by the National Human Genome Research Institute, National Institutes of Health (R01HG006753 to T.L.).

## Author contributions

A.H. and P.C. designed and developed tDRnamer standalone software. P.C. designed and developed tDRnamer web server. P.C. and T.L. wrote the manuscript. All authors contributed to the design of tDR naming convention and the feature requirements of tDRnamer.

## Competing interests

The authors declare no competing interests.

## Notes

http://trna.ucsc.edu/tDRnamer/

http://trna.ucsc.edu/tDRnamer/data/

### Competing Interest Statement

The authors have declared no competing interest.

