## Supplemental Tables and Figures for "A Standardized Ontology for Naming tRNA-derived RNAs Based on Molecular Origin"

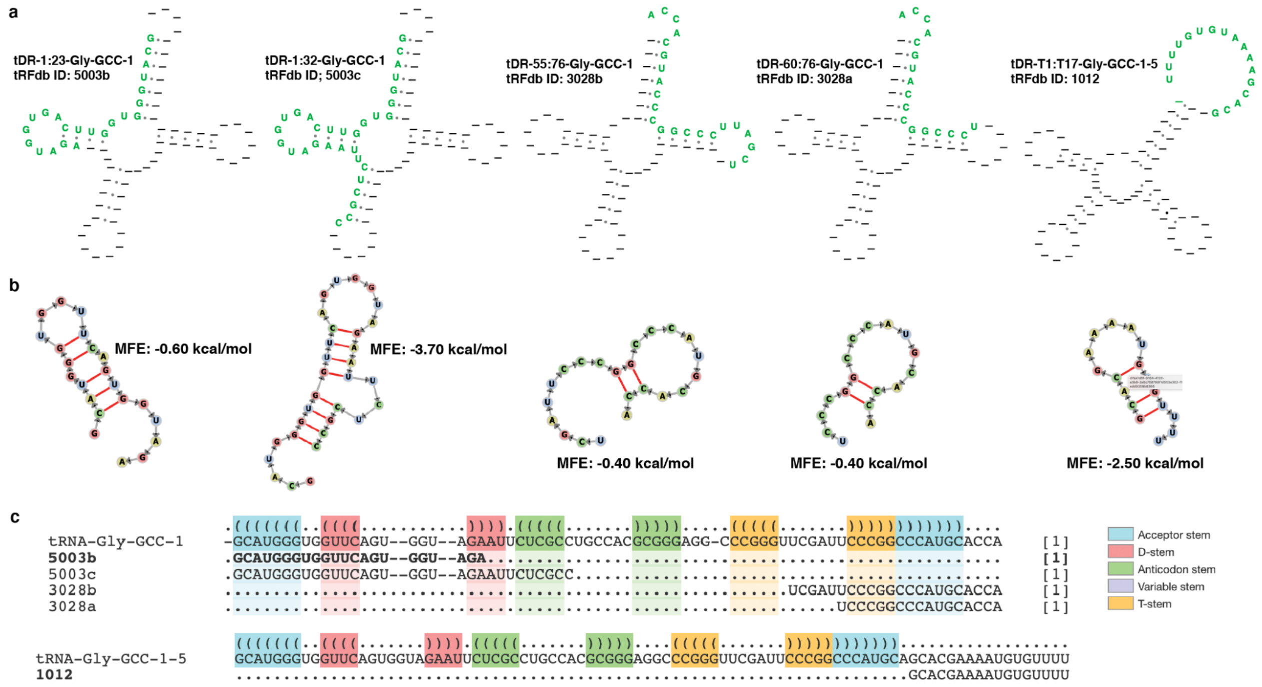


**Supplementary Fig. 1** A family of five tDRs derived from human tRNA-Gly-GCC-1. **a.** tDRs (two from 5′ end of the source tRNA, two from 3′ end, and one from 3′ trailer) are superimposed onto the secondary structure of tRNA-Gly-GCC-1. **b**. Minimum free energy secondary structures are predicted for tDRs. **c**. Four of the tDRs (tRFdb IDs: 5003b, 5003c, 3028a, and 3028b) are aligned with the mature tRNA-Gly-GCC-1 transcript. tDR-T1:T17-Gly-GCC-1-5 (tRFdb ID: 1012) is aligned separately to the precursor tRNA-Gly-GCC-1-5. All images are generated by the web server version of tDRnamer.

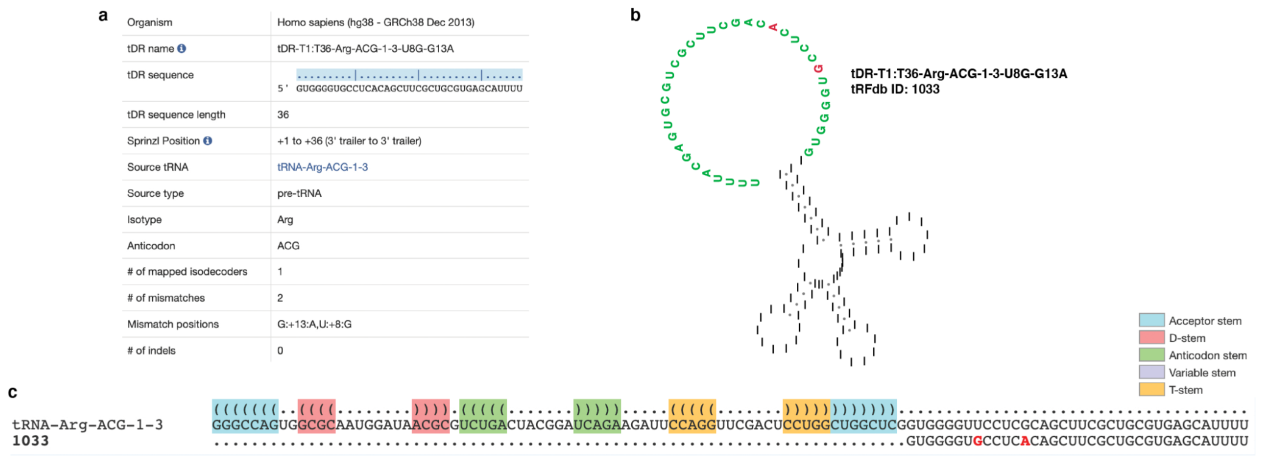


**Supplementary Fig. 2** tDR-T1:T36-Arg-ACG-1-3-U8G-G13A with two single nucleotide variations from 3′ trailer of tRNA$^{Arg(ACG)}$. **a.** tDR details are listed in the screenshot of tDRnamer results. **b**. tDR is superimposed onto the secondary structure of tRNA-Arg-ACG-1-3. **c**. tDR is aligned with source tRNA-Arg-ACG-1-3, showing known substitution mutations (dbSNP IDs: rs4982711 and rs34026312) marked in red. All images are generated by the web server version of tDRnamer.

**Supplementary Table 1** tDRnamer feature comparison in standalone and web server versions. Both versions employ the same processing routines for naming tDR sequences and searching tDR sequences by name. Feature differences in usage are listed.

| **Features** | **Standalone version** | **Web server version** |
|---|---|---|
| Use environment | Command-line in Linux/Unix platform | Web browser |
| Reference database | Download pre-built databases for model organisms or build user-defined databases with create_tDRnamer_db | Use available pre-built databases for model organisms (can request new databases to be added) |
| Input for tDR naming | Pre-processed small RNA sequencing reads in FASTQ file or tDR sequences in FASTA file | tDR sequences in FASTA format (maximum 1,000) |
| Input for tDR sequence search | Text file with tDR names | tDR names (maximum 500) |
| tDR sequence output | FASTA file | FASTA file and web page visualization |
| tDR naming and annotation output | Tab-delimited file | Tab-delimited file and web page visualization with links to source tRNAs in GtRNAdb |
| tDR alignment output | Stockholm files | Image visualization and downloadable files (PNG and postscript) |
| tDR secondary structure prediction | Not available | Minimum free energy (MFE) secondary structure image visualization and downloadable files (PNG and postscript) |
| tDR group output | Text file with alignments | Text file with alignments and web page visualization with links to tDR group members and source tRNAs in GtRNAdb |

**Supplementary Table 2** Re-annotation of tDRs derived from human tRNA$^{Pro}$ in tRFdb. Prefixes of License Plates in parentheses were assigned automatically when using the web server tool and depend on the existence of tDRs in MINTbase.

| **tRFdb ID** | **tDR position** | **tDR names and synonyms by tDRnamer** | **License Plates nomenclature** |
|---|---|---|---|
| 5011a | 1 to 15 | tDR-1:15-Pro-AGG-1-M8<br>tDR-1:15-Pro-AGG-2-M8<br>tDR-1:15-Pro-AGG-3-M8<br>tDR-1:15-Pro-CGG-1-M8<br>tDR-1:15-Pro-CGG-2-M8<br>tDR-1:15-Pro-TGG-1-M8<br>tDR-1:15-Pro-TGG-2-M8<br>tDR-1:15-Pro-TGG-3-M8 | (nlr-)15-6978WP |
| 5012a | 1 to 18 | tDR-1:18-Pro-AGG-1-M7<br>tDR-1:18-Pro-AGG-2-M7<br>tDR-1:18-Pro-AGG-3-M7<br>tDR-1:18-Pro-CGG-1-M7<br>tDR-1:18-Pro-CGG-2-M7<br>tDR-1:18-Pro-TGG-2-M7<br>tDR-1:18-Pro-TGG-3-M7 | (tRF-)17-6978WP4 |
| 5013b | 1 to 22 | tDR-1:22-Pro-AGG-1-M6<br>tDR-1:22-Pro-AGG-2-M6<br>tDR-1:22-Pro-CGG-1-M6<br>tDR-1:22-Pro-CGG-2-M6<br>tDR-1:22-Pro-TGG-2-M6<br>tDR-1:22-Pro-TGG-3-M6 | (tRF-)21-6978WPRLE |
| 5012b | 1 to 24 | tDR-1:24-Pro-AGG-1-M6<br>tDR-1:24-Pro-AGG-2-M6<br>tDR-1:24-Pro-CGG-1-M6<br>tDR-1:24-Pro-CGG-2-M6<br>tDR-1:24-Pro-TGG-2-M6<br>tDR-1:24-Pro-TGG-3-M6 | (tRF-)23-6978WPRL0L |
| 5013c | 1 to 31 | tDR-1:31-Pro-AGG-1-M5<br>tDR-1:31-Pro-AGG-2-M5<br>tDR-1:31-Pro-CGG-1-M5<br>tDR-1:31-Pro-CGG-2-M5<br>tDR-1:31-Pro-TGG-3-M5 | (tRF-)30-6978WPRLXN4V |
| 3002a | 59 to 76 | tDR-59:76-Pro-AGG-1-M8<br>tDR-59:76-Pro-AGG-2-M8<br>tDR-59:76-Pro-AGG-3-M8<br>tDR-59:76-Pro-CGG-1-M8<br>tDR-59:76-Pro-CGG-2-M8<br>tDR-59:76-Pro-TGG-1-M8 | (tRF-)18-HR6HFRD2 |

| tRFdb ID | tDR position | tDR names and synonyms by tDRnamer | License Plates nomenclature |
|---|---|---|---|
| | | tDR-59:76-Pro-TGG-2-M8<br>tDR-59:76-Pro-TGG-3-M8 | |
| 3002b | 55 to 76 | tDR-55:76-Pro-AGG-1-M8<br>tDR-55:76-Pro-AGG-2-M8<br>tDR-55:76-Pro-AGG-3-M8<br>tDR-55:76-Pro-CGG-1-M8<br>tDR-55:76-Pro-CGG-2-M8<br>tDR-55:76-Pro-TGG-1-M8<br>tDR-55:76-Pro-TGG-2-M8<br>tDR-55:76-Pro-TGG-3-M8 | (tRF-)22-8B8SOUPR2 |

**Supplementary Table 3** Files included in tDRnamer reference database. “dbname” in the file name represents the database name set as input option.

| File | Description |
|---|---|
| dbname-tRNAgenome.* | FASTA file of tRNA sequences with Bowtie2 indexes |
| dbname-trnaalign.stk | Alignments of mature tRNA sequences in Stockholm file format |
| dbname-trnaconvert.stk | Alignments of mature tRNA sequences in Stockholm file format |
| dbname-trnaloci.stk | Alignment of tRNA gene sequences in Stockholm file format |
| dbname-trnatable.txt | Tab-delimited file with tRNA transcripts and tRNA genes map |
| dbname-maturetRNAs.fa | FASTA file of mature tRNA sequences |
| dbname-maturetRNAs.bed | tRNA transcripts in BED file format |
| dbname-tRNAloci.fa | FASTA file of tRNA gene sequences |
| dbname-trnaloci.bed | tRNA genes in BED file format |
| dbname-filtered-tRNAs.out | tRNAscan-SE output file format with filtered tRNA genes used for database creation |
| dbname-dbinfo.txt | Database creation information |
| dbname-create_tDRnamer_db.log | Database creation log file |

**Supplementary Table 4** Output files generated by tDRnamer. “prefix” in the file name represents the output file prefix set by the --output option. “tDR” as prefix of file name represents the tDR name of the corresponding transcript.

| Output File | Description |
|---|---|
| prefix-tDR.fa | FASTA file with tDR names and sequences |
| prefix-tDR-info.txt | Tab-delimited file with tDR annotations including tDR names and sequences, source tRNAs, Sprinzl positions of tDRs relative to source tRNAs, original tRNA isotype and anticodon, sequence variation counts, and group ID |
| prefix-tDR-groups.txt | Text file containing queried tDRs that are grouped together. Alignments of group members and source tRNAs are arranged in Stockholm format that includes primary sequence and secondary structure information. |
| prefix-found-seq.fa | FASTA file with tDR sequences when searching by tDR names. This file is generated during the initial round of sequence search before the annotation process. |
| prefix-tDRs.stk | Alignments of identified tDRs derived from mature tRNAs with reference tRNA sequences in Stockholm file format |
| prefix-pre-tDRs.stk | Alignments of identified tDRs derived from precursor tRNAs with reference tRNA sequences in Stockholm file format. This file is only generated when the sequence source is set as eukaryotes. |
| prefix-unique-seq.fa | FASTA file with unique sequences in provided FASTQ file. This is only generated when a FASTQ file is provided as input. |
| prefix-filtered-seq.fa | FASTA file with sequences after filtering by length constraint and error checking. This is only generated when naming tDRs. |
| prefix-reformatted-seq.fa | FASTA file with sequences after converting RNA sequences to DNA sequences if applicable |
| prefix-filtered-names.txt | List of tDR names after filtering by error checking. This is only generated when searching by tDR names. |
| prefix-tDR-list.txt | Intermediate file generated during tDR annotation process |
| prefix-clusters.txt | Intermediate file generated during tDR group process |
| prefix-pre-clusters.txt | Intermediate file generated during tDR group process. This file is only generated when the sequence source is set as eukaryotes. |
| prefix-find-tdrs.log | Intermediate log file for searching tDR sequences by names |
| prefix_tDRnamer.log | Log file of tDRnamer run |

| **Output File** | **Description** |
|---|---|
| tDR-align.png | PNG file of tDR superimposed on source tRNA |
| tDR-align.ps | Postscript file of tDR superimposed on source tRNA |
| tDR-MFE.png | Secondary structure prediction of tDR based on minimum free energy in PNG file |
| tDR-MFE.eps | Secondary structure prediction of tDR based on minimum free energy in postscript file |
